# Verbal paired associates and the hippocampus: The role of scenes

**DOI:** 10.1101/206250

**Authors:** Ian A. Clark, Misun Kim, Eleanor A. Maguire

## Abstract

It is widely agreed that patients with bilateral hippocampal damage are impaired at binding pairs of words together. Consequently, the verbal paired associates (VPA) task has become emblematic of hippocampal function. This VPA deficit is not well understood, and is particularly difficult for hippocampal theories with a visuospatial bias to explain (e.g., cognitive map and scene construction theories). Resolving the tension among hippocampal theories concerning the VPA could be important for leveraging a fuller understanding of hippocampal function. Notably, VPA tasks typically use high imagery concrete words and so conflate imagery and binding. To determine why VPA engages the hippocampus, we devised an fMRI encoding task involving closely matched pairs of scene words, pairs of object words and pairs of very low imagery abstract words. We found that the anterior hippocampus was engaged during processing of both scene and object word pairs in comparison to abstract word pairs, despite binding occurring in all conditions. This was also the case when just subsequently remembered stimuli were considered. Moreover, for object word pairs, fMRI activity patterns in anterior hippocampus were more similar to those for scene imagery than object imagery. This was especially evident in participants who were high imagery users, and not in mid and low imagery users. Overall, our results show that hippocampal engagement during VPA, even when object word pairs are involved, seems to be evoked by scene imagery rather than binding. This may help to resolve the issue that visuospatial hippocampal theories have in accounting for verbal memory.

## INTRODUCTION

The field of hippocampal neuroscience is characterized by vigorous debates. But one point on which there is general agreement is that people with bilateral hippocampal damage and concomitant amnesia (hippocampal amnesia) are significantly impaired on verbal paired associates (VPA) tasks. The VPA task is a widely-used instrument for testing verbal memory and has been a continuous sub-test within the Wechsler Memory Scale from its initial inception (Wechsler, 1945) to the present day (WMS-IV; Wechsler, 2009). While the VPA task has been revised many times (e.g., increasing the number of word pairs to be remembered, changing the ratio of difficult to easy word pairs) the basic premise has remained the same. The requirement is to encode pairs of words (e.g., bag–truck), memory for which is then tested. Testing can be conducted in multiple ways, but one primary outcome measure is performance on a delayed cued recall test (i.e. the experimenter asks for the word that goes with bag) 30 minutes after the completion of the learning trials. Compared to matched healthy control participants, patients with hippocampal amnesia show a consistent and reliable deficit on delayed cued recall tests (Giovanello, Verfaellie, & Keane, 2003; Graf & Schacter, 1985; Spiers, Maguire, & Burgess, 2001; Zola-Morgan, Squire, & Amaral, 1986), and consequently the VPA has become emblematic of hippocampal function.

The VPA task is typically regarded as a verbal memory task. However, many theories focus on elucidating the role of the hippocampus in visuospatial rather than verbal processing. This includes accounts that consider spatial navigation (Maguire et al., 2000; O’Keefe & Nadel, 1978), autobiographical memory (Hassabis & Maguire, 2007; Scoville & Milner, 1957; Squire, 1992), scene perception (Graham, Barense, & Lee, 2010; McCormick, Rosenthal, Miller, & Maguire, 2017), the mental construction of visual scene imagery (Maguire & Mullally, 2013; Zeidman & Maguire, 2016) and more specific aspects of visuospatial processing, including perceptual richness, a sense of reliving and imagery content (Andrews-Hanna, Reidler, Sepulcre, Poulin, & Buckner, 2010; St-Laurent, Moscovitch, & McAndrews, 2016; St. Jacques, Conway, Lowder, & Cabeza, 2010).

The cognitive map theory, for instance, posits that the hippocampus specifically supports flexible, allocentric representations of spatial relationships (O’Keefe & Nadel, 1978). In contrast, the scene construction theory (see also the emergent memory account; Graham et al., 2010) proposes that the anterior hippocampus constructs models of the world in the form of spatially coherent scenes (Dalton & Maguire, 2017; Hassabis & Maguire, 2007; Maguire & Mullally, 2013; Zeidman & Maguire, 2016). A scene in this context is a specific type of visual image that represents a naturalistic three-dimensional space typically populated by objects and that is viewed from an egocentric perspective. The construction of scene imagery involves associative processing and binding, but the scene construction theory asserts that the hippocampus is specifically required to perform these functions in the service of creating scene representations (Maguire & Mullally, 2013). The difficulty with theories such as cognitive map and scene construction is that they do not appear to be able to explain why VPA learning is invariably compromised following hippocampal damage.

On the face of it, another hippocampal theory does seem to account for the VPA findings. The relational theory suggests that the hippocampus makes associations between any elements, regardless of whether or not space or scenes are involved (Cohen & Eichenbaum, 1993; Konkel & Cohen, 2009). This generic associative process could account for the creation of an association between two unrelated words in the VPA task, while also explaining the involvement of the hippocampus in visuospatial tasks, and the combining of individual elements into a coherent memory, or the recombination of different elements from past experiences to simulate the future (Moscovitch, Cabeza, Winocur, & Nadel, 2016; Roberts, Schacter, & Addis, 2017; Schacter et al., 2012; St. Jacques, Carpenter, Szpunar, & Schacter, 2018; Thakral, Benoit, & Schacter, 2017). However, a purely associative account of hippocampal function is not completely satisfactory given that patients with hippocampal damage retain an ability to form associations in some circumstances (see Clark & Maguire, 2016).

Resolving the tension among hippocampal theories concerning the VPA could be important for leveraging a fuller understanding of hippocampal function. In taking this issue forwards, it is worthwhile first to step back. Examination of the words used in typical VPA tests shows the vast majority are high imagery concrete words. It could be that people use visual imagery when processing the word pairs (Maguire & Mullally, 2013). This speculation has recently received indirect support from the finding that patients with hippocampal amnesia used significantly fewer high imagery words in their narrative descriptions of real and imagined events (Hilverman, Cook, & Duff, 2017), suggesting a potential link between verbal processing and visual imagery.

Currently, therefore, standardised VPA tests may be conflating associative processes and imageability. Patients with hippocampal damage are reportedly unable to imagine fictitious and future scenes in addition to their well reported memory deficits (Hassabis, Kumaran, Vann, & Maguire, 2007; Race, Keane, & Verfaellie, 2011; Schacter et al., 2012). It would, therefore, follow that their impoverished scene imagery ability may place them at a disadvantage for processing high imagery concrete words. One way to deal with the conflation of visual imagery and binding is to examine very low imagery (abstract) word pairs, which would assess binding outside of the realm of imagery. However, abstract word pairs rarely feature in VPA tests used with patients or in neuroimaging experiments.

In addition, different types of high imagery words are not distinguished in VPA tests, with the majority of words representing single objects. However, the scene construction theory links the anterior hippocampus specifically with constructing visual imagery of scenes (Dalton & Maguire, 2017; Zeidman & Maguire, 2016). By contrast, the processing of single objects is usually associated with perirhinal and lateral occipital cortices (Malach et al., 1995; Murray, Bussey, & Saksida, 2007). It could therefore be that a scene word (e.g., forest) in a pair engages the hippocampus (via scene imagery) and not because of binding or visual imagery in general. It has also been suggested that even where each word in a pair denotes an object (e.g., cat–table), this might elicit imagery of both objects together in a scene and it is the generation of this scene imagery that recruits the hippocampus (Clark & Maguire, 2016; Maguire & Mullally, 2013). Consequently, if visual imagery does play a role in the hippocampal-dependence of the VPA task, then it will be important to establish not only whether visual imagery or binding is more relevant, but also the type of visual imagery being employed.

To determine why VPA engages the hippocampus, we devised an fMRI task with three types of word pairs: where both words in a pair denoted Scenes, where both words represented single Objects, and where both words were very low imagery Abstract words. This allowed us to separate imageability from binding, and to examine different types of imagery. Of particular interest were the Object word pairs because we wanted to ascertain whether they were processed using scene or object imagery. For all word pairs, our main interest was during their initial presentation, when any imagery would likely be evoked.

In addition, we conducted recognition memory tests after scanning to investigate whether the patterns of (hippocampal) activity were affected by whether pairs were successfully encoded or not. While the VPA memory test used with patients typically involves cued recall, the adaptation of the VPA task for fMRI necessitated the use of recognition memory tests. This is because performing a cued recall test for 135 word pairs that were each seen only once is too difficult even for healthy participants. Finally, given that people vary in their use of mental imagery (Kosslyn, Brunn, Cave, & Wallach, 1984; Marks, 1973; McAvinue & Robertson, 2007), we also tested groups of high, mid and low imagery users to assess whether this influenced hippocampal engagement during VPA encoding.

In line with the scene construction theory, we hypothesised that anterior hippocampal activity would be apparent for Scene words pairs, given the likely evocation of scene imagery. We also predicted that anterior hippocampal activity would be increased for Object word pairs, and that this would be best explained by the use of scene imagery. In addition, we expected that the effect of scene imagery use on the hippocampus would be most apparent in high imagery users. By contrast, we predicted that Abstract words pairs would engage areas outside of the hippocampus, even when only subsequently-remembered pairs were considered.

## METHODS

### Participants

Forty five participants took part in the fMRI study. All were healthy, right-handed, and had normal or corrected to normal vision. Given the verbal nature of the task, all participants were highly proficient in English, had English as their first language and were educated in English throughout their school years. Each participant gave written informed consent. The study was approved by the University College London Research Ethics Committee. Participants were recruited on the basis of their scores on the Vividness of Visual Imagery Questionnaire (VVIQ; Marks, 1973). The VVIQ is a widely-used self-report questionnaire which asks participants to bring images to mind and rate them on a 5 point scale as to their vividness (anchored at 1: “perfectly clear and as vivid as normal vision”, and 5: “No image at all, you only ‘know’ that you are thinking of the object”). Therefore, a high score on the VVIQ corresponds to low use of visual imagery. The validity of the VVIQ has been demonstrated in numerous ways. For example, experimental studies have found that high visualisers were able to match two pictures more quickly than low visualisers when the first picture had to be retained as a mental image over a 20 second period (Gur & Hilgard, 1975). Additionally, significant correlations between the VVIQ and the Betts’ Questionnaire Upon Mental Imagery (another widely-used imagery questionnaire, Sheehan, 1967) have also been reported (Burton & Fogarty, 2003; Campos & Pérez-Fabello, 2005).

Our fMRI participants comprised three sub-groups (n=15 in each), low imagery users, mid imagery users and high imagery users. Initially, 184 people completed the VVIQ. Fifteen of the highest and 15 of the lowest scorers made up the low and high imagery groups. A further 15 mid scorers served as the mid imagery group. We acknowledge that these groups are relatively small for an fMRI study, but we were nevertheless interested to see whether any differences would be observed. The groups did not differ significantly on age, gender, years of education and general intellect. Table 1 provides details of the three groups.

**Table 1.** Characteristics of the participant groups.

|  | Imagery Group |  |  | p-value |  |  |
| --- | --- | --- | --- | --- | --- | --- |
|  | Low | Mid | High | Low vs Mid | Low vs High | Mid vs High |
| Age | 23.07 (2.31) | 21.87 (2.20) | 23.93 (5.26) | 0.16 | 0.57 | 0.18 |
| Number of males | 6 (40.0%) | 7 (46.67%) | 8 (53.33%) | 0.71 | 0.46 | 0.72 |
| Years of education | 16.0 (1.89) | 15.8 (1.61) | 16.0 (2.33) | 0.76 | 1.0 | 0.79 |
| Matrix Reasoning | 12.47 (2.26) | 11.47 (2.17) | 12.07 (3.61) | 0.23 | 0.72 | 0.59 |
| TOPF | 54.93 (5.13) | 57.47 (5.49) | 53.0 (9.47) | 0.20 | 0.49 | 0.13 |
| FSIQ | 110.13 (5.48) | 111.97 (6.13) | 110.22 (5.99) | 0.39 | 0.97 | 0.44 |
| VCI | 108.93 (5.32) | 110.81 (6.18) | 108.75 (6.0) | 0.38 | 0.93 | 0.36 |
| VVIQ mean score | 3.08 (0.45) | 2.15 (0.17) | 1.51 (0.25) | < 0.001 | < 0.001 | < 0.001 |

Means (standard deviations). Two-tailed p-values for t-tests ( $\chi^2$ test for the number of males). General intellect was measured using the Matrix Reasoning subtest (scaled scores) of the Wechsler Adult Intelligence Scale-IV (WAIS-IV; Wechsler, 2008) and the Test of Premorbid Function (TOPF; Wechsler, 2011) providing an estimate of Full Scale IQ (FSIQ) and a Verbal Comprehension Index (VCI). VVIQ=Vividness of Visual Imagery Questionnaire.

### Stimuli

To ensure that any fMRI differences were due to our imagery manipulation and not other word properties, the word conditions were highly matched. Six hundred and fifty four words were required for the study – 218 Scene words, 218 Object words and 218 Abstract words. Words were initially sourced from databases created by Brysbaert and colleagues, which provided ratings for concreteness, word frequency, age of acquisition, valence and arousal (Brysbaert, Warriner, & Kuperman, 2014; Kuperman, Stadthagen-Gonzalez, & Brysbaert, 2012; van Heuven, Mandera, Keuleers, & Brysbaert, 2014; Warriner, Kuperman, & Brysbaert, 2013). It was important to control for valence and arousal given reports of higher emotional ratings for abstract words, which could influence fMRI activity (Kousta, Vigliocco, Vinson, Andrews, & Del Campo, 2011; Vigliocco et al., 2014). We also used data from the English Lexicon Project (Balota et al., 2007) to provide lexical information about each word – word length, number of phonemes, number of syllables, number of orthographic neighbours, and number of phonological and phonographic neighbours with and without homophones.

To verify that each word induced the expected imagery (i.e., scene imagery, object imagery or very little/no imagery for the abstract words), we collected two further ratings for each word. First, a rating of imageability to ensure that Scene and Object words were not only concrete but also highly imageable (although concreteness and imageability are often interchanged, and while they are highly related constructs, they are not the same; Paivio, Yuille, & Madigan, 1968), and additionally that Abstract words were low on imageability. Second, a decision was elicited about the type of imagery the word brought to mind. This was in response to the following instruction: “If you had an image we would like you to classify it as either a ‘scene’ or an ‘object’. A scene is an image in your mind that has a sense of space; that you could step into or operate within. An object on the other hand is more of an isolated image, without additional background imagery. It is also likely that for a number of words you will experience very little or no imagery – please do select this option if this is the case”. These ratings were collected from 119 participants in total using Amazon Mechanical Turk’s crowdsourcing website, following the procedures employed by Brysbaert and colleagues for the databases described above. Words were classified as a Scene or Object word when there was a minimum of 70% agreement on the type of imagery brought to mind, and the mean imageability rating was greater than 3.5 (out of 5). For Abstract words, the mean imageability had to be less than or equal to 2. An overview of the word properties is shown in Table 2. This also includes summary comparison statistics. A list of the words in each category can be found at:
*http://www.fil.ion.ucl.ac.uk/Maguire/Clark_et_al_2018_Scene_Words.pdf*
*http://www.fil.ion.ucl.ac.uk/Maguire_Clark_et_al_2018_Object_Words.pdf*
*http://www.fil.ion.ucl.ac.uk/Maguire/Clark_et_al_2018_Abstract_Words.pdf*

**Table 2.** Properties of each word type.

| Word property | Type of word |  |  | p-value |  |  |
| --- | --- | --- | --- | --- | --- | --- |
|  | Scene | Object | Abstract | Scene vs Object | Scene vs Abstract | Object vs Abstract |
| **Lexical criteria** |  |  |  |  |  |  |
| Number of letters<sup>a</sup> | 6.73 (1.99) | 6.64 (1.91) | 6.75 (1.97) | 0.64 | 0.90 | 0.55 |
| Number of phonemes<sup>a</sup> | 5.61 (1.89) | 5.41 (1.71) | 5.66 (1.72) | 0.25 | 0.77 | 0.13 |
| Number of syllables<sup>a</sup> | 2.09 (0.85) | 2.06 (0.79) | 2.16 (0.77) | 0.68 | 0.38 | 0.18 |
| Number of orthographic neighbours<sup>a</sup> | 2.57 (4.97) | 3.09 (4.96) | 2.35 (4.33) | 0.28 | 0.62 | 0.10 |
| Number of phonological neighbours<sup>a</sup> | 6.28 (11.71) | 7.61 (11.80) | 6.12 (11.31) | 0.23 | 0.89 | 0.18 |
| Number of phonological neighbours (inc homophones)<sup>a</sup> | 6.88 (12.59) | 8.13 (12.48) | 6.66 (11.86) | 0.30 | 0.85 | 0.21 |
| Number of phonographic neighbours<sup>a</sup> | 1.53 (3.51) | 1.78 (3.48) | 1.46 (3.21) | 0.45 | 0.84 | 0.32 |
| Number of phonographic neighbours (inc homophones)<sup>a</sup> | 1.61 (3.67) | 1.96 (3.63) | 1.49 (3.36) | 0.32 | 0.72 | 0.16 |
| Word frequency: Zipf<sup>b</sup> | 3.90 (0.71) | 3.80 (0.61) | 3.88 (0.82) | 0.12 | 0.77 | 0.27 |
| Age of acquisition<sup>c</sup> | 7.69 (2.14) | 7.40 (2.12) | 9.78 (2.46) | 0.15 | < 0.001 | < 0.001 |
| **Emotional constructs** |  |  |  |  |  |  |
| Valence<sup>d</sup> | 5.68 (1.08) | 5.63 (1.02) | 5.58 (1.12) | 0.63 | 0.34 | 0.61 |
| Number of positive words<sup>d,e</sup> | 171 (78.44%) | 173 (79.6%) | 167 (76.61%) | 0.81 | 0.65 | 0.49 |
| Hedonic valence<sup>d,f</sup> | 1.07 (0.69) | 0.98 (0.68) | 1.04 (0.70) | 0.18 | 0.69 | 0.34 |
| Arousal<sup>d</sup> | 4.07 (0.96) | 3.99 (0.87) | 4.04 (0.71) | 0.34 | 0.73 | 0.46 |
| **Imagery** |  |  |  |  |  |  |
| Concreteness<sup>g</sup> | 4.65 (0.22) | 4.68 (0.22) | 1.83 (0.29) | 0.11 | < 0.001 | < 0.001 |
| Imageability<sup>h</sup> | 4.38 (0.29) | 4.41 (0.32) | 1.53 (0.20) | 0.31 | < 0.001 | < 0.001 |

Means (standard deviations). Two-tailed p-values for t-tests ( $\chi^2$ test for the number of positive words). Note that each comparison was assessed separately in order to provide a greater opportunity for any differences between conditions to be identified.
<sup>a</sup>From the English Lexicon Project (Balota et al., 2007: exlexicon.wustl.edu).
<sup>b</sup>From van Heuven et al. (2014). The Zipf scale is a standardised measure of word frequency using a logarithmic scale. Values go from 1 (low frequency words) to 6 (high frequency words).
<sup>c</sup>From Kuperman et al. (2012).
<sup>d</sup>From Warriner et al. (2013).
<sup>e</sup>Positive words were those that had a valence score greater than or equal to 5.
<sup>f</sup>Hedonic valence is the distance from neutrality (i.e., from 5), regardless of being positive or negative, as per Vigliocco et al. (2014).
<sup>g</sup>From Brysbaert et al. (2014).
<sup>h</sup>Collected for the current study as detailed in the Methods.

Scene, Object and Abstract words were matched on 13 out of the 16 measures. Scene and Object words were matched on all 16 measures, whereas Abstract words, as expected, were less concrete and less imageable than Scene and Object words and had a higher age of acquisition, as is normal for abstract words (Kuperman et al., 2012; Stadthagen-Gonzalez & Davis, 2006). As well as being matched at the overall word type level as shown on Table 2, within each word type, words were assigned to one of four lists (word pairs, single words, catch trials or post-scan memory test lures), and all lists were matched on all measures.

### Experimental design and task

The fMRI task consisted of two elements, the main task and catch trials. The latter were included to provide an active response element and to encourage concentration during the experiment. To match the WMS-IV Verbal Paired Associate Test (Wechsler, 2009), each stimulus was presented for 4 seconds. This was followed by a jittered baseline (a central fixation cross) for between 2 and 5 seconds which aided concentration by reducing the predictability of stimulus presentation (Figure 1D). The scanning session was split into four runs, three runs containing 80 trials lasting 10 minutes each and a final run of 78 trials lasting 9 minutes 45 seconds. Trials were presented randomly for each participant with no restrictions on what could precede or follow each trial.

**Figure 1.**
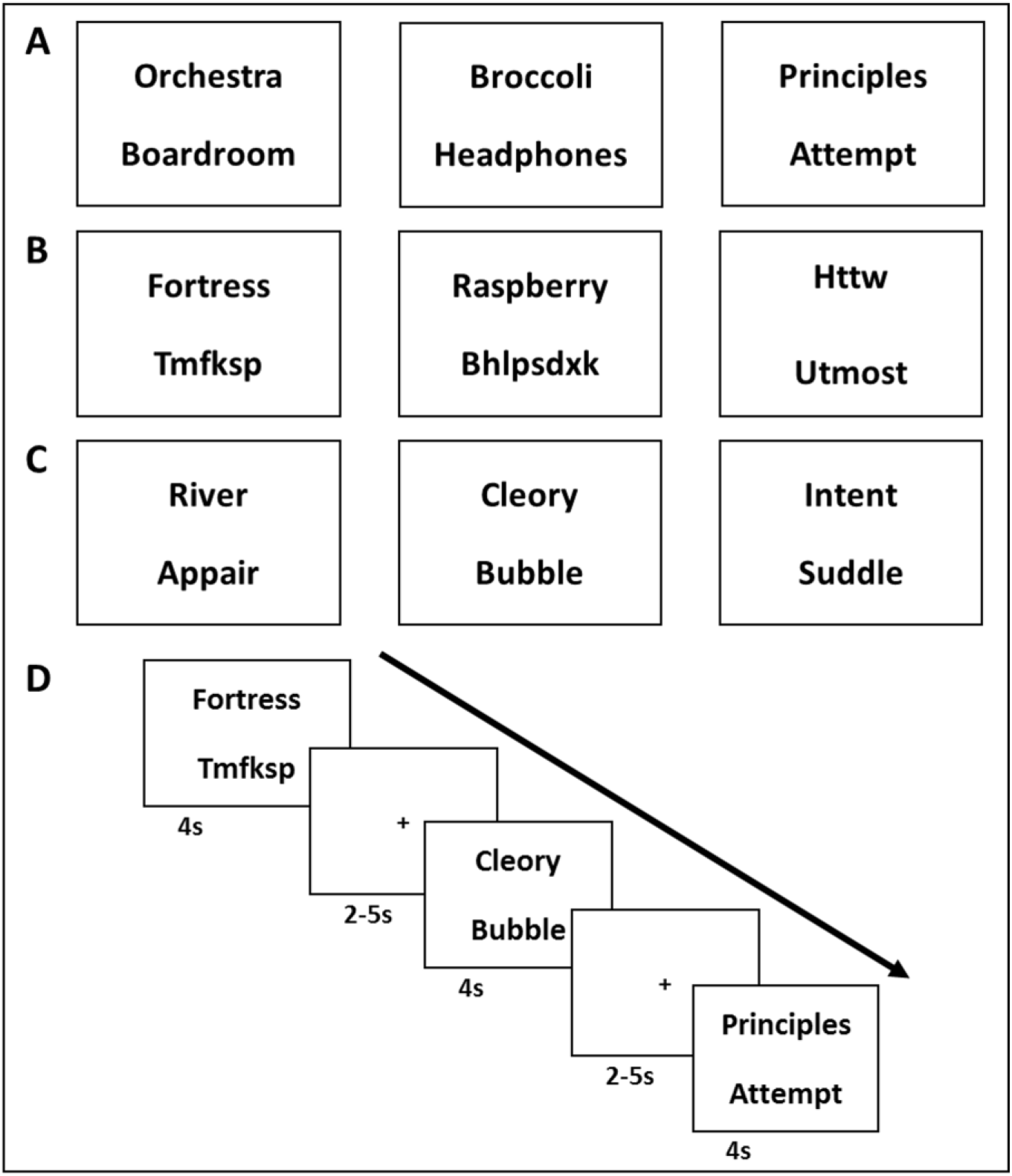
Example stimuli and trial timeline. **(A)** Examples of stimuli from each of the word types in the order of (from left to right) Scene word pair, Object word pair, Abstract word pair. **(B)** Examples of single word trials in the order of (from left to right) Scene single word, Object single word, Abstract single word. Single words were shown with random letter strings (which could be presented at either the top or the bottom) in order to be similar to the visual presentation of the word pairs. **(C)** Examples of catch trials, where a real word was presented with a pseudoword, which could be presented as either the top or bottom word. **(D)** Example timeline of several trials.

Unbeknownst to participants, there were six categories of stimuli – high imagery Scene words, high imagery Object words and very low imagery Abstract words, shown either in pairs of the same word type (Figure 1A) or as single words (Figure 1B). To equalise visual presentation between the word pairs and the single words, the latter were presented with a random letter string that did not follow the rules of the English language and did not resemble real words (Figure 1B). The average, minimum and maximum length of the letter strings was matched to the real words. Letter strings could either be presented at the top or the bottom of the screen. There were 45 trials of each condition, with each word shown only once to the participant. Our prime interest was in the word pair conditions, and in particular the Object word pairs, as these related directly to our research question. The single word conditions were included for the purposes of specific analyses, which are detailed in the Results section.

Participants were asked to try and commit the real words to memory for later memory tests, and were specifically instructed that they would be asked to recall the pairs of real words as pairs. They were explicitly told they would not need to remember the random letter strings. No further instructions about how to memorise the stimuli were given (i.e., we did not tell participants to use any particular strategy).

Participants were told that occasionally there would be catch trials where they had to indicate, using a button press, if they saw a real word presented with a ‘pseudoword’ (Figure 1C). The participants were informed that they were not required to remember the real word or the pseudoword presented in these catch trials. A pseudoword is a combination of letters that resembles a real English word and follows the rules of the English language, but is not an actual real word. Pseudowords were generated using the English Lexicon Project (Balota et al., 2007) and were paired with Scene, Object or Abstract words. They were presented at either the top or the bottom of the screen to ensure that participants attended to both. The number of letters and orthographic neighbours of the pseudowords were matched to all of the real word conditions and across the three pseudoword groups (all p’s > 0.3). Additionally, across the pseudoword groups we matched the accuracy of pseudoword identification (all p’s > 0.6) as reported in the English Lexicon Project (Balota et al., 2007). Forty eight catch trials were presented over the course of the experiment, 16 trials with each of the word types, ranging between 10 and 15 in each of the four runs. Catch trials were pseudo-randomly presented to ensure regular presentation but not in a predictable manner. Feedback was provided at the end of each scanning run as to the number of correctly identified pseudowords and incorrectly identified real words.

### Post-scan recognition memory tests

Following scanning, participants had two recognition memory tests. The first was an item recognition memory test for all 405 words presented during scanning (45 words for each of three single word types, and 90 words for each of three paired word types) and a further 201 foils (67 of each word type). Each word was presented on its own in the centre of the screen for up to 5 seconds. Words were presented randomly in a different order for each participant. Participants had to indicate for each word whether they had seen it in the scanner (old) or not (new). Following this, they rated their confidence in their answer on a 3 point scale – high confidence, low confidence or guessing. Any trials where a participant correctly responded “old” and then indicated they were guessing were excluded from subsequent analyses.

After the item memory test, memory for the pairs of words was examined. This associative memory test presented all of the 135 word pairs shown to participants in the scanner and an additional 66 lure pairs (22 of each type), one pair at a time, for up to 5 seconds. The word pairs were presented in a different random order for each participant. The lure pairs were constructed from the single words that were presented to the participants in the scanner. Therefore, the participants had seen all of the words presented to them in the associative recognition memory test, but not all were previously in pairs, specifically testing whether the participants could remember the correct associations. Participants were asked to indicate whether they saw that exact word pair presented to them in the scanner (old) or not (new). They were explicitly told that some pairs would be constructed from the single words they had seen during scanning and not to make judgements solely on individual words, but to consider the pair itself. Confidence ratings were obtained in the same way as for the item memory test, and trials where a participant correctly responded “old” and then indicated they were guessing were excluded from subsequent analyses.

### Debriefing

On completion of the memory tests, participants were asked about their strategies for processing the words while they were in the scanner. At this point, the participants were told about the three different types of words presented to them – Scenes, Objects and Abstract. For each word type, and separately for single words and word pairs, participants were presented with reminders of the words. They were then asked to choose from a list of options as to which strategy best reflected how they processed that word type. Options included: “I had a visual image of a scene related to this type of single word” (scene imagery), “I had a visual image of a single entity (e.g., one specific object) for a word with no other background imagery” (object imagery), “I read each word without forming any visual imagery at all” (no imagery).

### Statistical analyses of the behavioural data

#### Stimuli creation and participant group comparisons

Comparisons between word conditions, and between the participant groups, were performed using independent samples t-tests for continuous variables and chi squared tests for categorical variables. An alpha level of p > 0.05 was used to determine that the stimuli/groups were matched. Note that each comparison was assessed separately (using t-tests or chi squared tests) in order to provide a greater opportunity for any differences between conditions to be identified.

#### Main study

Both within and between participants designs were used. The majority of analyses followed a within-participants design, with all participants seeing all word conditions. Additionally, participants were split into three groups dependent on their VVIQ score allowing for between-participants analyses to be performed.

All data were assessed for outliers, defined as values that were at least 2 standard deviations away from the mean. If an outlier was identified then the participant was removed from the analysis in question (and this is explicitly noted in the Results section). Memory performance for each word condition was compared to chance level (50%) using one sample t-tests. For all within-participants analyses, when comparing across three conditions, repeated measures ANOVAs with follow-up paired t-tests were employed, and for comparison across two conditions paired t-tests were utilised. For between-participants analyses a one-way ANOVA was performed with follow up independent samples t-tests.

All ANOVAs were subjected to Greenhouse-Geisser adjustment to the degrees of freedom if Mauchly’s sphericity test identified that sphericity had been violated. For all statistical tests alpha was set at 0.05. Effect sizes are reported following significant results as Cohen’s d for one sample and independent sample t-tests, Eta squared for repeated measures ANOVA and Cohen’s d for repeated measures (d_rm_) for paired samples t-tests (Lakens, 2013). All analyses were performed in IBM SPSS statistics v22.

### Scanning parameters and data pre-processing

T2*-weighted echo planar images (EPI) were acquired using a 3T Siemens Trio scanner (Siemens Healthcare, Erlangen, Germany) with a 32-channel head coil. fMRI data were acquired over four scanning runs using scanning parameters optimised for reducing susceptibility-induced signal loss in the medial temporal lobe: 48 transversal slices angled at −30°, TR=3.36 s, TE=30 ms, resolution=3 × 3 × 3mm, matrix size=64 × 74, z-shim gradient moment of −0.4mT/m ms (Weiskopf, Hutton, Josephs, & Deichmann, 2006). Fieldmaps were acquired with a standard manufacturer’s double echo gradient echo field map sequence (short TE=10 ms, long TE=12.46 ms, 64 axial slices with 2 mm thickness and 1 mm gap yielding whole brain coverage; in-plane resolution 3 × 3 mm). After the functional scans, a 3D MDEFT structural scan was obtained with 1mm isotropic resolution (Deichmann, Schwarzbauer, & Turner, 2004).

Preprocessing of data was performed using SPM12 (www.fil.ion.ucl.ac.uk/spm). The output of the SPM image realignment protocol showed that head motion was low (mean (SD) in mm: X = 0.51 (0.32), Y = 1.29 (0.33), Z = 1.64 (0.80); mean (SD) in degrees: pitch = 0.03 (0.03), Roll = 0.01 (0.01), Yaw = 0.01 (0.01)), and was smaller than the voxel size. Functional images were co-registered to the structural image, and then realigned and unwarped using field maps. The participant’s structural image was segmented and spatially normalised to a standard EPI template in MNI space with a voxel size of 2 × 2 × 2mm and the normalisation parameters were then applied to the functional data. For the univariate analyses, the functional data were smoothed using an 8mm full-width-half-maximum Gaussian kernel. In line with published RSA literature (e.g. Chadwick, Jolly, Amos, Hassabis, & Spiers, 2015; Kriegeskorte, Mur, Ruff, et al., 2008; Marchette, Vass, Ryan, & Epstein, 2014), the multivariate analyses used unsmoothed data. We used unsmoothed data in order to capture neural information in the form of spatially distributed activity across multiple voxels. Smoothing potentially washes out the fine activity differences between voxels.

Where bilateral region of interest (ROI) analyses were performed, the hippocampal ROIs were manually delineated on a previously collected (n = 36) group averaged structural MRI scan (1 × 1 × 1 mm) using ITK-SNAP (www.itksnap.org) and then resampled to our functional scans (2 × 2 × 2 mm). The anterior hippocampus was delineated using an anatomical mask that was defined in the coronal plane and went from the first slice where the hippocampus can be observed in its most anterior extent until the final slice of the uncus. In terms of structural space, this amounted to 3616 voxels and in functional space to 481 voxels. The posterior hippocampus was defined as proceeding from the first slice following the uncus until the final slice of observation in its most posterior extent (see Dalton, Zeidman, Barry, Williams, & Maguire, 2017 for more details). In terms of structural space, this amounted to 4779 voxels and in functional space to 575. The whole hippocampus mask combined the anterior and posterior masks, and therefore contained 8395 voxels in structural space and 1056 voxels in functional space.

### fMRI analysis: univariate

The six experimental word conditions were Scene, Object and Abstract words, presented as either word pairs or single words. As noted above, our prime interest was in the word pair conditions, and in particular the Object word pairs, as these related directly to our research question. We therefore directly contrasted fMRI BOLD responses between the word pair conditions. The single word conditions were included for the purposes of specific analyses, which are detailed in the Results section. We performed two types of whole brain analysis, one using all of the trials (45 per condition) and another using only trials where the items were subsequently remembered, not including trials where the participant indicated they were guessing. The average number of trials per condition were: Scene word pairs 31.49 (SD = 6.25); Object word pairs 34.53 (SD = 5.84); Abstract word pairs 27.42 (SD = 8.67). See Table 4 for comparisons of the number of correct trials, not including guessing, across the conditions.

For both analyses, the GLM consisted of the word condition regressors convolved with the haemodynamic response function, in addition to participant-specific movement regressors and physiological noise regressors. The Artifact Detection Toolbox (http://www.nitrc.org/projects/artifact_detect/) was used to identify spikes in global brain activation and these were entered as a separate regressor. Participant-specific parameter estimates for each regressor of interest were calculated for each voxel. Second level random effects analyses were then performed using one sample t-tests on the parameter estimates. For comparison across VVIQ imagery groups, we performed an ANOVA with follow-up independent sample t-tests. We report results at a peak-level threshold of p less than 0.001 whole-brain uncorrected for our a priori region of interest – the hippocampus – and p less than 0.05 family-wise error (FWE) corrected at the voxel-level elsewhere.

In addition, several ROI analyses were performed on a subset of the univariate analyses. Three ROIs were considered – the whole hippocampus, the anterior hippocampus and the posterior hippocampus (all bilateral). We used a peak-level threshold of p less than 0.05 FWE-corrected at the voxel level for each mask and, where indicated in the Results section, also a more lenient threshold of p less than 0.001 uncorrected for each mask.

### fMRI analysis: multivariate

Multivoxel pattern analysis was used to test whether the neural representations of the Object word pairs were more similar to the Scene single words than the Object single words when separately examining bilateral anterior and posterior hippocampal ROIs. For each participant, T-statistics for each voxel in the ROI were computed for each condition (Object word pair, Object single word, Scene single word) and in each scanning run. The Pearson correlation between each condition was then calculated as a similarity measure (Object word pair/Object word pair, Object word pair/Scene single word, Object word pair/Object single word). The similarity measure was cross-validated across the different scanning runs to guarantee the independence of each data set. Repeated measures ANOVA and paired t-tests were used to compare the similarity between conditions at the group level. This multivariate analysis was first applied to the data from all participants, and then to the three subsets of participants (low, mid and high imagery users). All data were assessed for outliers, defined as values that were at least 2 standard deviations away from the group mean. If an outlier was identified then the participant was removed from the analysis in question (and this is explicitly noted in the Results section).

Note that the absolute correlation of the similarity value is expected to be low due to inherent neural variability and the fact that a unique set of words was presented for each scanning run. As such, the important measure is the comparison of the similarity value between the conditions, not the absolute similarity value of a single condition. The range of similarity values that we found was entirely consistent with those reported in other studies employing a similar representational similarity approach in a variety of learning, memory and navigation tasks in a wide range of brain regions (Bellmund, Deuker, Navarro Schröder, & Doeller, 2016; Chadwick et al., 2015; Deuker, Bellmund, Navarro Schröder, & Doeller, 2016; Hsieh, Gruber, Jenkins, & Ranganath, 2014; Hsieh & Ranganath, 2015; Kim, Jeffery, & Maguire, 2017; Milivojevic, Vicente-Grabovetsky, & Doeller, 2015; Schapiro, Turk-Browne, Norman, & Botvinick, 2016; Schuck, Cai, Wilson, & Niv, 2016; Staresina, Henson, Kriegeskorte, & Alink, 2012).

## Results

### Behavioural

On average, participants identified 85.56% (SD=11.52) of the pseudowords during catch trials, showing that they maintained concentration during the fMRI experiment. On the post-scan item memory test, Scene, Object and Abstract words were remembered above chance and there were no differences between the conditions (Table 3, which includes the statistics). Performance on the associative memory test also showed that Scene, Object and Abstract word pairs were remembered above chance (Table 4, which includes the statistics). Considering the average performance across the four word conditions used in the main univariate analyses (i.e. Scene word pairs, Object word pairs, Abstract word pairs, Abstract single words) then one participant performed below chance. The fMRI analyses do not change whether this participant is included or not. Comparison of memory performance across the word types found differences in performance in line with the literature (Paivio, 1969). Both types of high imagery word pairs (Scene and Object) were remembered better than Abstract word pairs (Figure 2; Table 4), while Object word pairs were remembered better than Scene word pairs. Given that the word pair memory lures were highly confusable with the actual word pairs (because the lure pairs were made up of the studied single words), d’ values were also calculated for the word pairs. Scene, Object and Abstract word pairs all showed d’ values greater than 0, representing the ability to discriminate between old and new pairs (Table 4). Both Scene and Object word pairs had greater d’ values than Abstract word pairs, and Object word pairs d’ values were greater than those for Scene word pairs (Table 4), showing the same pattern as that calculated using the percentage correct. Overall, these behavioural findings show that, despite the challenging nature of the experiment with so many stimuli, participants engaged with the task, committed a good deal of information to memory and could successfully distinguish between previously-presented word pairs and highly confusable lures.

**Figure 2.**
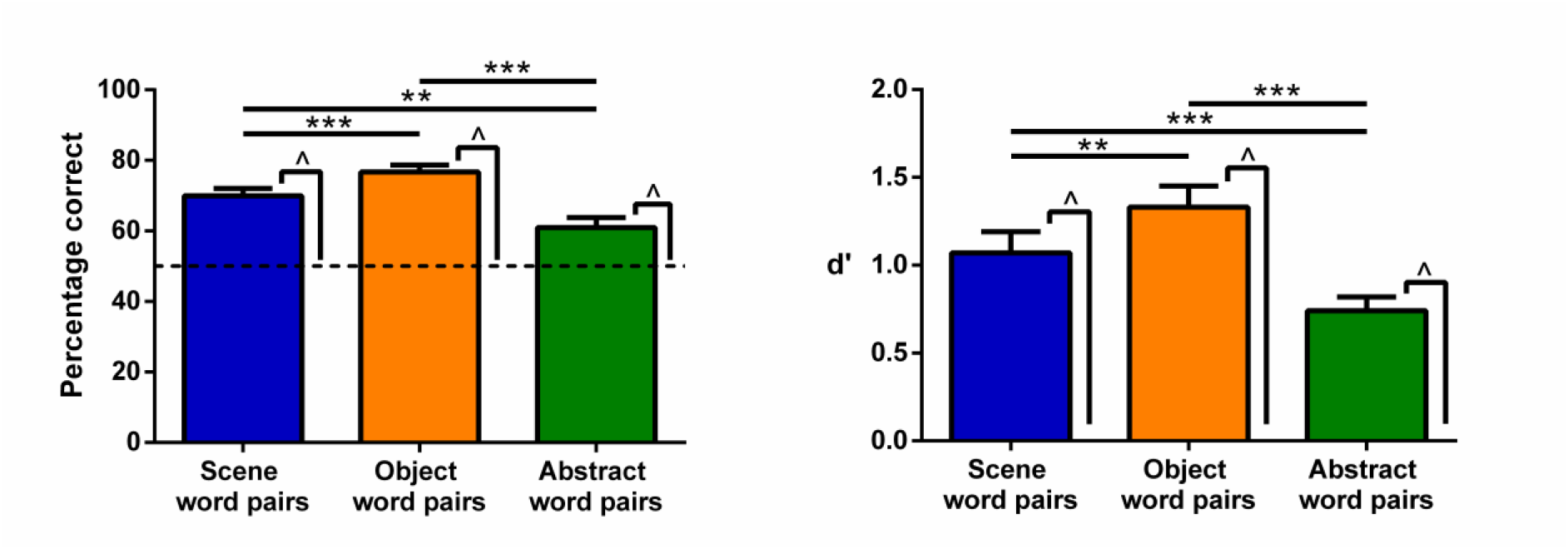
Memory performance on the associative memory test shown by percentage correct (left) and d’ (right). Error bars are 1 standard error of the mean. ^ indicates a significant difference from chance (for percentage correct the dashed line indicates chance at 50%, for d’ it is 0) at p < 0.001. Stars show the significant differences across the word pair types; **p < 0.01, *** p < 0.001.

**Table 3.** Performance (% correct) on the post-scan item memory test (non-guessing trials).

|  | Scene<br>single words | Object<br>single words | Abstract<br>single words |
| --- | --- | --- | --- |
| Mean | 67.41 | 66.37 | 67.61 |
| Standard Deviation | 14.93 | 17.71 | 16.06 |
| Comparison to chance (50%) |  |  |  |
|  | **t**<sub>44</sub> | **p** | **d** |
| Scene single words | 7.82 | < 0.001 | 2.36 |
| Object single words | 6.20 | < 0.001 | 1.87 |
| Abstract single words | 7.36 | < 0.001 | 2.22 |
| Comparison across the word types |  |  |  |
|  | **F**<sub>1.76,77.51</sub> | **p** |  |
| Main effect | 0.28 | 0.73 |  |

**Table 4.**
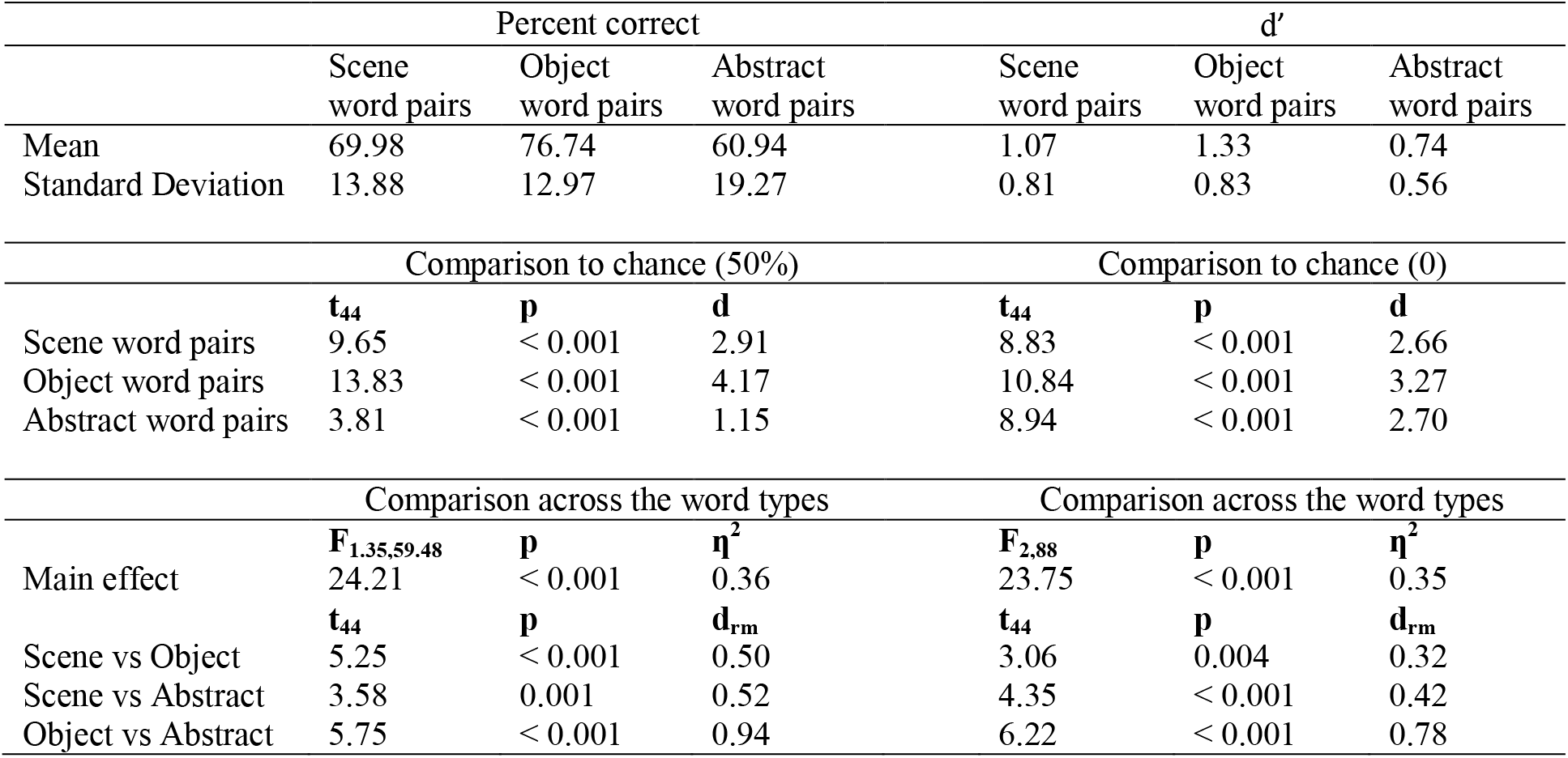
Performance (% correct and d’) on the post-scan associative memory test (non-guessing trials).

### fMRI

#### Univariate analyses

We performed two whole brain analyses, one using all of the trials and another using only trials where the items were subsequently remembered in the post-scan memory tests (the item memory test for the single word trials, the associative memory test for the word pairs, excluding trials where participants correctly responded “old” and then indicated they were guessing). The two analyses yielded very similar results across the whole brain, even though the analysis using only subsequently-remembered stimuli was less well powered due to the reduced number of stimuli. Given that our interest was in the point at which participants were initially processing the word pairs and potentially using mental imagery to do so, we focus on the results of the analysis using all of the trials. Results are also reported for the analyses using only the remembered stimuli, which allowed us to control for any memory-related effects.

We first compared the high imagery (Scene, Object) and very low imagery (Abstract) word pairs. All of the conditions involved associative processing, and so we reasoned that any differences we observed, particularly in hippocampal engagement, would be due to the imageability of the Scene and Object word pairs. As predicted, Scene word pairs elicited greater bilateral anterior (but not posterior) hippocampal activity compared to Abstract word pairs (Figure 3A, full details in Table 5A). Of note, increased activity was also observed in bilateral parahippocampal, fusiform, retrosplenial and left ventromedial prefrontal (vmPFC) cortices. The analysis using only the remembered stimuli showed very similar results, including for the anterior hippocampus (Table 6A). The reverse contrast identified no hippocampal engagement, but rather greater activity for Abstract word pairs in middle temporal cortex (−58, −36, −2, T=6.58) and temporal pole (−52, 10, - 22, T=6.16).

**Figure 3.**
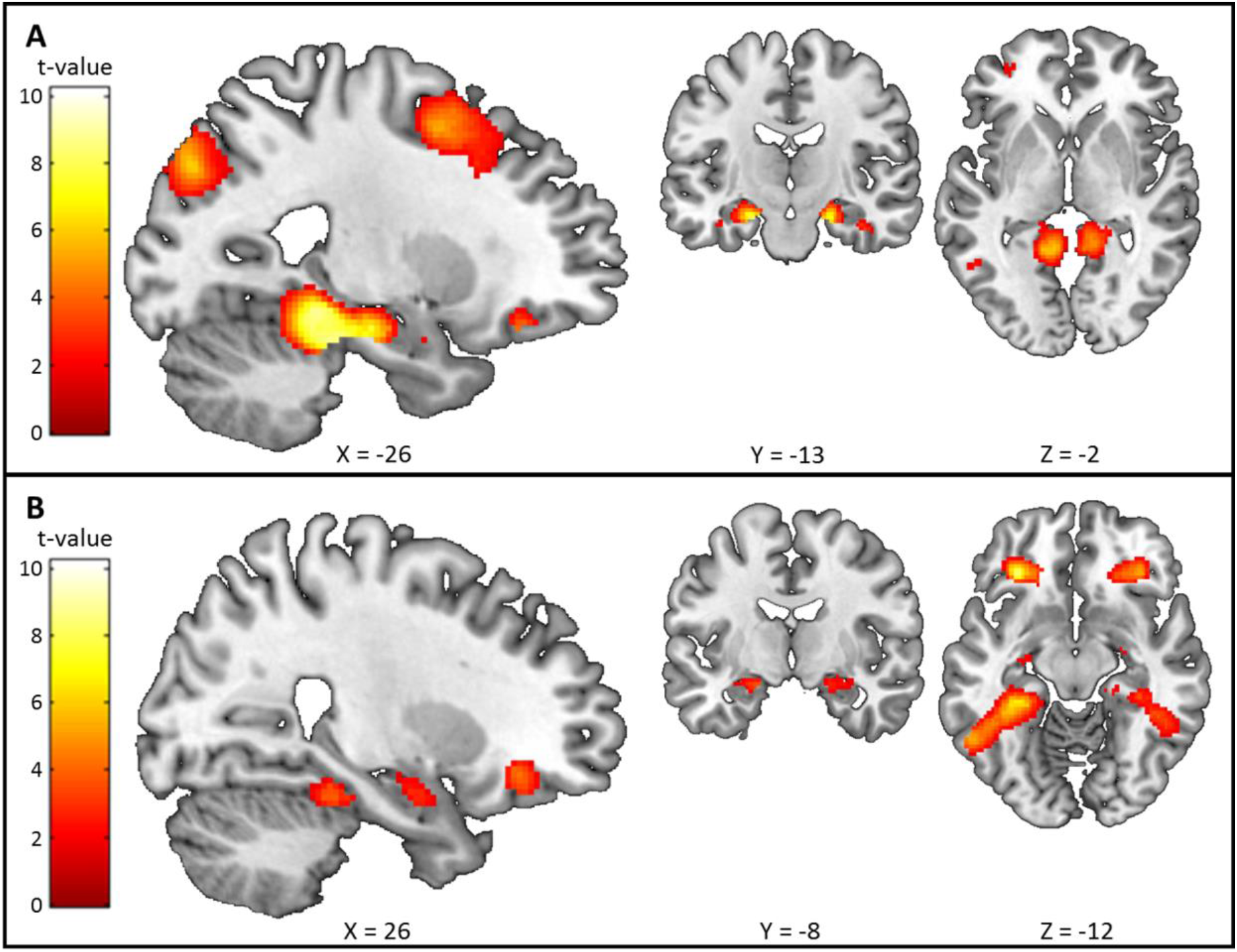
Comparison of imageable Scene or Object word pairs with non-imageable Abstract word pairs. The sagittal slice is of the left hemisphere which is from the ch2better template brain in MRicron (Holmes et al., 1998; Rorden & Brett, 2000). The left of the image is the left side of the brain. The coloured bar indicates the t-value associated with each voxel. **(A)** Scene word pairs > Abstract word pairs. **(B)** Object word pairs > Abstract word pairs. Images are thresholded at p < 0.001 uncorrected for display purposes.

**Table 5.**
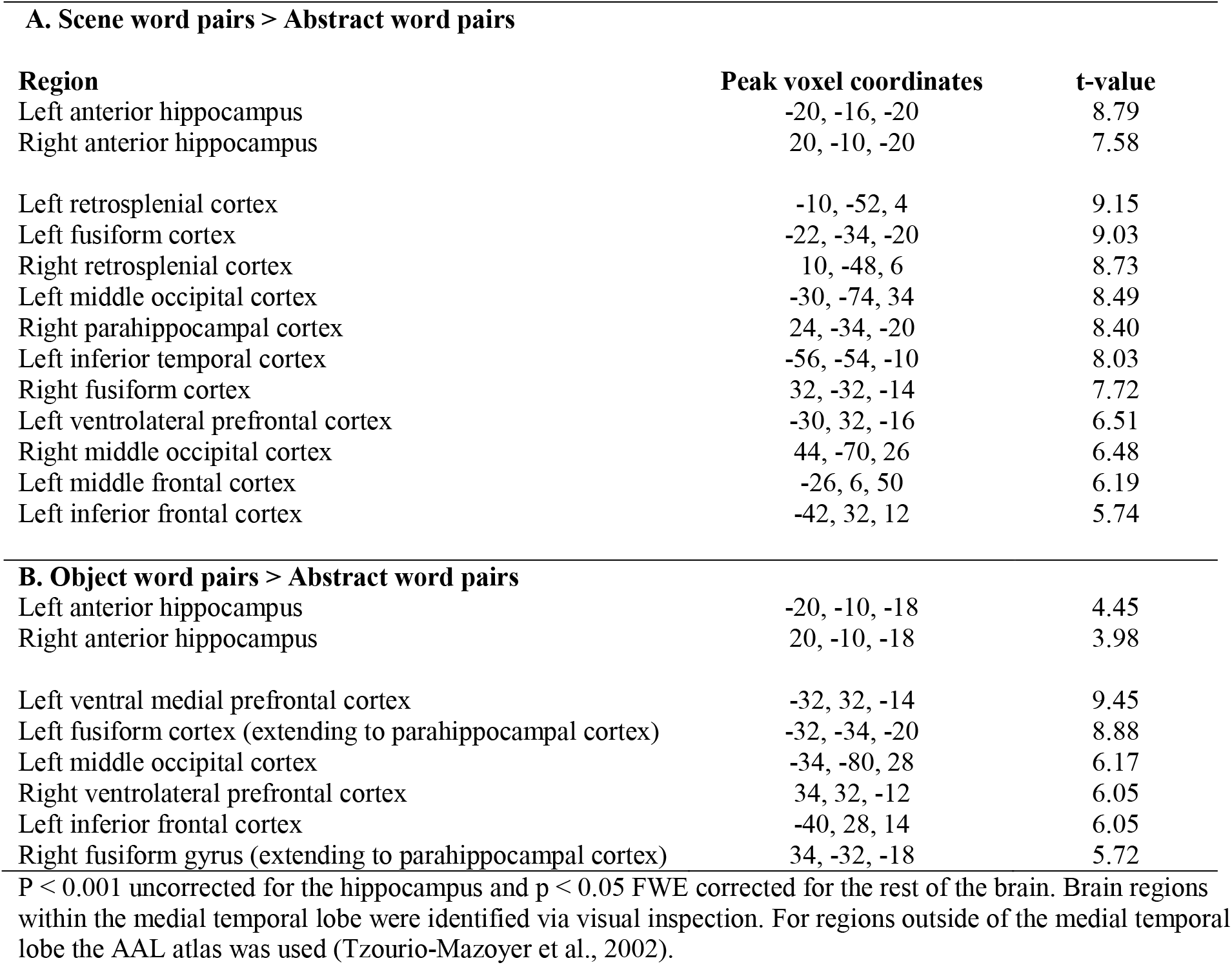
Imageable word pairs compared with Abstract word pairs.

| **A. Scene word pairs > Abstract word pairs** |  |  |
| --- | --- | --- |
| **Region** | **Peak voxel coordinates** | **t-value** |
| Left anterior hippocampus | -20, -16, -20 | 8.79 |
| Right anterior hippocampus | 20, -10, -20 | 7.58 |
| Left retrosplenial cortex | -10, -52, 4 | 9.15 |
| Left fusiform cortex | -22, -34, -20 | 9.03 |
| Right retrosplenial cortex | 10, -48, 6 | 8.73 |
| Left middle occipital cortex | -30, -74, 34 | 8.49 |
| Right parahippocampal cortex | 24, -34, -20 | 8.40 |
| Left inferior temporal cortex | -56, -54, -10 | 8.03 |
| Right fusiform cortex | 32, -32, -14 | 7.72 |
| Left ventrolateral prefrontal cortex | -30, 32, -16 | 6.51 |
| Right middle occipital cortex | 44, -70, 26 | 6.48 |
| Left middle frontal cortex | -26, 6, 50 | 6.19 |
| Left inferior frontal cortex | -42, 32, 12 | 5.74 |
| **B. Object word pairs > Abstract word pairs** |  |  |
| Left anterior hippocampus | -20, -10, -18 | 4.45 |
| Right anterior hippocampus | 20, -10, -18 | 3.98 |
| Left ventral medial prefrontal cortex | -32, 32, -14 | 9.45 |
| Left fusiform cortex (extending to parahippocampal cortex) | -32, -34, -20 | 8.88 |
| Left middle occipital cortex | -34, -80, 28 | 6.17 |
| Right ventrolateral prefrontal cortex | 34, 32, -12 | 6.05 |
| Left inferior frontal cortex | -40, 28, 14 | 6.05 |
| Right fusiform gyrus (extending to parahippocampal cortex) | 34, -32, -18 | 5.72 |
| P < 0.001 uncorrected for the hippocampus and p < 0.05 FWE corrected for the rest of the brain. Brain regions within the medial temporal lobe were identified via visual inspection. For regions outside of the medial temporal lobe the AAL atlas was used (Tzourio-Mazoyer et al., 2002). |  |  |

Object word pairs also showed greater bilateral anterior (but not posterior) hippocampal activity compared with the Abstract word pairs, along with engagement of bilateral parahippocampal cortex, fusiform cortex and vmPFC (Figure 3B, Table 5B), with increased anterior hippocampal activity also apparent when just the subsequently-remembered stimuli were considered (Table 6B). The reverse contrast identified no hippocampal engagement, but rather greater activity for Abstract word pairs in middle temporal cortex (−62, −32, −2, T=8) and temporal pole (−54, 10, −18, T=7.12).

**Table 6.** Remembered imageable word pairs compared with remembered Abstract word pairs.

| **A. Scene word pairs remembered > Abstract word pairs remembered** |  |  |
| --- | --- | --- |
| **Region** | **Peak voxel coordinates** | **T value** |
| Left hippocampus | -28, -22, -18 | 7.53 |
| Right hippocampus | 24, -20, -18 | 5.09 |
| Left retrosplenial cortex | -10, -50, 2 | 10.20 |
| Left fusiform gyrus (extending to parahippocampal gyrus) | -30, -34, -14 | 8.21 |
| Right retrosplenial cortex | 10, -48, 4 | 7.47 |
| Left middle occipital lobe | -30, -80, 40 | 7.23 |
| Right fusiform gyrus (extending to parahippocampal gyrus) | 26, -28, -20 | 7.00 |
| Left ventral medial prefrontal cortex | -30, 34, -12 | 6.51 |
| Right middle occipital lobe | 44, -70, 28 | 6.29 |
| Left inferior temporal gyrus | -56, -54, -10 | 5.64 |
| **B. Object word pairs remembered > Abstract word pairs remembered** |  |  |
| **Region** | **Peak voxel coordinates** | **T value** |
| Left hippocampus | -32, -22, -12 | 5.05 |
| Left ventral medial prefrontal cortex | -30, 34, -12 | 9.26 |
| Left fusiform gyrus (extending to parahippocampal gyrus) | -30, -32, -18 | 7.94 |
| Left middle occipital lobe | -34, -82, 30 | 6.43 |
| Left inferior temporal gyrus | -54, -58, -6 | 6.13 |

$P < 0.001$ uncorrected for the hippocampus and $p < 0.05$ FWE corrected for the rest of the brain. Brain regions within the medial temporal lobe were identified via visual inspection. For regions outside of the medial temporal lobe the AAL atlas was used (Tzourio-Mazoyer et al., 2002).

Increased anterior hippocampal activity was therefore observed for both Scene and Object word pairs compared to the very low imagery Abstract word pairs. As greater anterior hippocampal engagement was apparent even when using just the remembered stimuli, it is unlikely that this result can be explained by better associative memory or successful encoding for the high imagery word pairs. Rather the results suggest that the anterior hippocampal activity for word pair processing may be related to the use of visual imagery.

All of the above contrasts involved word pairs, suggesting that associative binding per se cannot explain the results. However, it could still be the case that binding Abstract word pairs does elicit increased hippocampal activity but at a lower level than Scene and Object word pairs. To address this point, we compared the Abstract word pairs with the Abstract single words, as this should reveal any hippocampal activity related to associative processing of the pairs. No hippocampal engagement was evident for the Abstract word pairs in comparison to the Abstract single words (Table 7). This was also the case when just the remembered stimuli were considered (Table 8), albeit with slightly lower power than the previous contrasts (number of trials for the Abstract word pairs = 27.42 (SD = 8.67); for the Abstract single words: 30.42 (SD = 7.23)).

**Table 7.** Abstract word pairs compared with Abstract single words.

| Abstract word pairs > Abstract single words |  |  |
| --- | --- | --- |
| Region | Peak voxel coordinates | t-value |
| Left middle temporal cortex | -64, -36, 2 | 8.39 |
| Left temporal pole | -52, 12, -16 | 6.72 |
| Left fusiform cortex | -38, -46, -20 | 6.64 |
| Left inferior frontal cortex | -54, 24, 12 | 6.54 |
| Left inferior occipital cortex | -42, -68, -12 | 6.52 |
| Right inferior occipital cortex | 36, -74, -12 | 6.11 |
| Right lingual cortex | 20, -82, -10 | 5.87 |
| Left pre-central gyrus | -50, 0, 48 | 5.84 |

P < 0.001 uncorrected for the hippocampus (no activations found) and p < 0.05 FWE corrected for the rest of the brain. Brain regions within the medial temporal lobe were identified via visual inspection. For regions outside of the medial temporal lobe the AAL atlas was used (Tzourio-Mazoyer et al., 2002).

**Table 8.** Remembered Abstract word pairs compared with remembered Abstract single words.

| **Abstract word pairs remembered > Abstract single words remembered** |  |  |
| --- | --- | --- |
| **Region** | **Peak voxel coordinates** | **T value** |
| Left inferior frontal gyrus | -54, 14, 12 | 9.50 |
| Left pre-central gyrus | -48, -2, 48 | 8.02 |
| Left middle temporal gyrus | -52, -46, 4 | 8.21 |
| Left inferior occipital lobe | -38, -78, -8 | 7.23 |
| Right inferior occipital lobe | 34, -80, -6 | 7.11 |
| Left supplementary motor area | -2, 4, 56 | 6.72 |
| Right inferior frontal gyrus | 50, 10, 28 | 6.44 |
| Right superior temporal pole | 46, -30, 4 | 6.11 |
| Right caudate nucleus | 12, 10, 6 | 6.07 |
| Left pallidum | -18, 6, 0 | 6.04 |

P < 0.001 uncorrected for the hippocampus (no activations found) and p < 0.05 FWE corrected for the rest of the brain. Brain regions within the medial temporal lobe were identified via visual inspection. For regions outside of the medial temporal lobe the AAL atlas was used (Tzourio-Mazoyer et al., 2002).

Given the difficulty of interpreting null results, in particular when using whole-brain standard contrasts, we performed additional ROI analyses to further test whether any sub-threshold hippocampal activity was evident for the Abstract word pairs compared to the Abstract single words. Using an anatomically-defined bilateral whole hippocampal mask, no differences in hippocampal activity were apparent at a p<0.05 FWE-corrected threshold at the voxel level for the mask, or when a more lenient p<0.001 uncorrected threshold was employed. We then extracted average beta values from across the whole hippocampus bilateral ROI, and two additional smaller ROIs – anterior and posterior hippocampus – for the Abstract word pairs and Abstract single words. T-tests showed that there were no differences between conditions (whole hippocampus: t_44_=0.16, p=0.88; anterior hippocampus only: t_44_=0.13, p=0.89; posterior hippocampus only: t_44_=0.18, p=0.86). Similar results were also observed when using just the remembered stimuli (whole hippocampus: t_44_=1.16, p=0.25; anterior hippocampus only: t_44_=1.36, p=0.18; posterior hippocampus only: t_44_=0.63, p=0.53). Overall, therefore, even at lenient thresholds and using an ROI approach, no hippocampal engagement was identified for Abstract word pairs compared to the Abstract single words. While the absence of evidence is not evidence of absence, this is in direct contrast to our findings of increased hippocampal activity for the high imagery word pairs compared to the very low imagery Abstract word pairs. This, therefore, lends support to the idea that the use of visual imagery might be important for inducing hippocampal responses to word pairs.

We also predicted that anterior hippocampal activity would be specifically influenced by the use of scene imagery, as opposed to visual imagery per se. The inclusion of both Scene and Object word pairs offered the opportunity to test this. Scene word pairs would be expected to consistently evoke scene imagery (as both words in a pair represented scenes), while Object word pairs could evoke both or either object and scene imagery (e.g., object imagery by imagining the two objects without a background context, or scene imagery by creating a scene and placing the two objects into it), thus potentially diluting the hippocampal scene effect. Scene word pairs might therefore activate the anterior hippocampus to a greater extent than Object word pairs. This comparison also provided an additional opportunity to contrast the effects of scene imagery and memory performance on hippocampal activity, because Object word pairs were better remembered than the Scene word pairs. As such, if hippocampal activity could be better explained by word pair memory performance rather than scene imagery, we would expect that Object word pairs would show greater hippocampal activity than Scene word pairs.

Contrasting Scene and Object word pairs revealed that, in line with our prediction, Scene word pairs evoked greater bilateral anterior (but not posterior) hippocampal activity than the Object word pairs (Figure 4, Table 9A). Analysis using just the remembered stimuli gave similar results (Table 10A). Other areas that showed increased activity for the Scene word pairs included the retrosplenial and parahippocampal cortices. The reverse contrast examining what was more activated for Object word pairs compared to Scene word pairs found no evidence of hippocampal activity despite better subsequent memory performance for the Object word pairs (Table 9B), even when just the remembered stimuli were examined (Table 10B). It seems, therefore, that the anterior hippocampus may be particularly responsive to scene imagery and that increases in hippocampal activity in this task were not driven by greater memory performance.

**Table 9.** Scene word pairs compared with Object word pairs.

| **A. Scene word pairs > Object word pairs** |  |  |
| --- | --- | --- |
| **Region** | **Peak voxel coordinates** | **t-value** |
| Left anterior hippocampus | -22, -18, -20 | 5.55 |
| Right anterior hippocampus | 22, -20, -20 | 6.07 |
| Right retrosplenial cortex | 16, -54, 20 | 7.35 |
| Left retrosplenial cortex | -10, -50, 4 | 7.34 |
| Left fusiform cortex (extending to parahippocampal cortex) | -28, -38, -12 | 7.25 |
| Right fusiform cortex (extending to parahippocampal cortex) | 28, -26, -20 | 6.87 |
| Left middle temporal cortex | -58, -6, -14 | 5.77 |
| **B. Object word pairs > Scene word pairs** |  |  |
| Left inferior temporal cortex | -42, -48, -16 | 7.16 |

P < 0.001 uncorrected for the hippocampus and p < 0.05 FWE corrected for the rest of the brain. Brain regions within the medial temporal lobe were identified via visual inspection. For regions outside of the medial temporal lobe the AAL atlas was used (Tzourio-Mazoyer et al., 2002).

**Table 10.** Remembered Scene word pairs compared with remembered Object word pairs.

| **A. Scene word pairs remembered > Object word pairs remembered** |  |  |
| --- | --- | --- |
| **Region** | **Peak voxel coordinates** | **T value** |
| Right hippocampus | 24, -20, -20 | 5.18 |
| Left hippocampus | -22, -20, -18 | 4.26 |
| Left retrosplenial cortex | -12, -50, 4 | 6.74 |
| Right fusiform gyrus (extending to parahippocampal gyrus) | 24, -28, -18 | 6.49 |
| Right retrosplenial cortex | 10, -48, 6 | 6.46 |
| Left fusiform gyrus (extending to parahippocampal gyrus) | -24, -38, -12 | 6.37 |
| **B. Object word pairs remembered > Scene word pairs remembered** |  |  |
| **Region** | **Peak voxel coordinates** | **T value** |
| Left inferior temporal gyrus | -42, -48, -16 | 6.12 |

P < 0.001 uncorrected for the hippocampus and p < 0.05 FWE corrected for the rest of the brain. Brain regions within the medial temporal lobe were identified via visual inspection. For regions outside of the medial temporal lobe the AAL atlas was used (Tzourio-Mazoyer et al., 2002).

**Figure 4.**
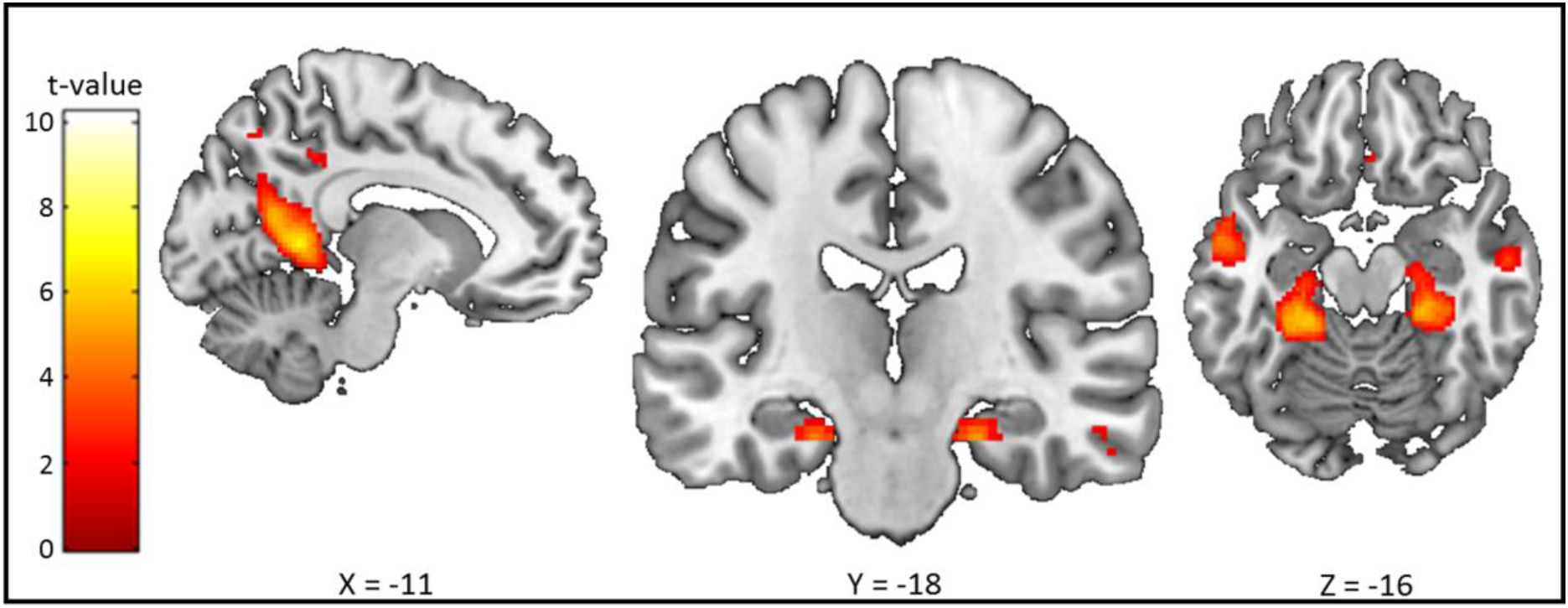
Brain areas more activated by Scene word pairs than Object word pairs. The sagittal slice is of the left hemisphere which is from the ch2better template brain in MRicron (Holmes et al., 1998; Rorden & Brett, 2000). The left of the image is the left side of the brain. The coloured bar indicates the t-value associated with each voxel. Images are thresholded at p < 0.001 uncorrected for display purposes.

To summarise, our univariate analyses found that Scene word pairs engaged the anterior hippocampus the most, followed by the Object word pairs, with the Abstract word pairs not eliciting any significant increase in activation (Figure 5). This is what we predicted, and may be suggestive of particular responsivity of the anterior hippocampus to scenes.

**Figure 5.**
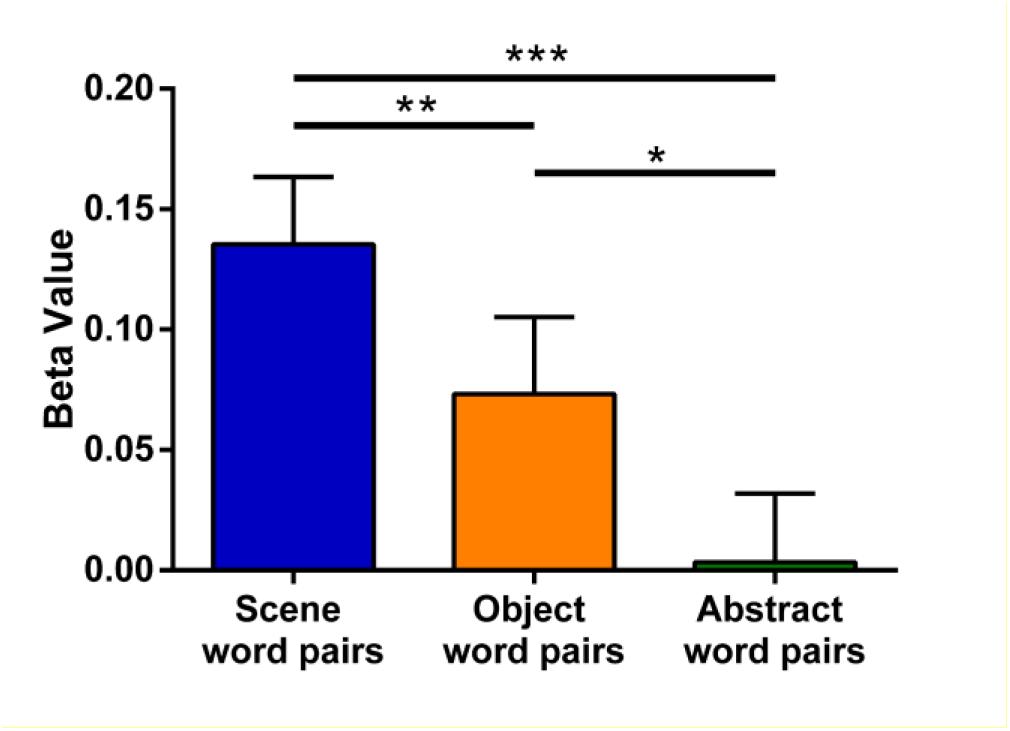
Comparison of each word pair condition with a fixation cross baseline. Mean beta values extracted from a bilateral anatomical mask of the anterior hippocampus for each of the word pair conditions compared to the central fixation cross baseline. Error bars are 1 standard error of the mean. A repeated measures ANOVA showed significant differences between the conditions (F_1,69,74.51_=16.06, p <0.001, η^2^=0.27). Follow-up paired t-tests revealed significant differences between Scene word pairs versus Abstract word pairs t_44_=6.46, p<0.001, d_rm_=0.70; Scene word pairs versus Object word pairs t_44_=2.97, p=0.005, d_rm_=0.30; Object word pairs versus Abstract word pairs t_44_= 2.51, p=0.016, drm=0.34. *p < 0.05, **p < 0.01, ***p<0.001.

#### Multivariate analyses

We next sought further, more direct, evidence that our main condition of interest, Object word pairs, elicited hippocampal activity via scene imagery. Given our univariate findings of increased anterior hippocampal activity for Scene word pairs and Object word pairs compared to Abstract word pairs, and the extant literature showing the importance of the anterior hippocampus for processing scenes (e.g. Zeidman & Maguire, 2016, but see also Sheldon & Levine, 2016), we looked separately at anatomically-defined bilateral anterior and posterior hippocampal ROIs. We then used multivariate representational similarity analysis (RSA; Kriegeskorte, Mur, & Bandettini, 2008) to compare the neural patterns of activity associated with encoding Object word pairs with Scene or Object single words. We predicted that the neural representations of Object word pairs in the anterior hippocampus would be more similar to Scene single words than Object single words, but that this would not be apparent in the posterior hippocampus. As our aim was to specifically investigate the contribution of different types of imagery to hippocampal activity, the scene and object single words were chosen as comparators because they consistently elicit either scene or object imagery respectively (see Methods). Abstract words do not elicit much visual imagery, so they were not included in the RSA analyses.

Three similarity correlations were calculated. First, the similarity between Object word pairs and themselves, which provided a baseline measure of similarity (i.e., the correlation of Object word pairs over the 4 runs of the scanning experiment). The two similarities of interest were the similarity between Object word pairs and Scene single words, and the similarity between Object word pairs and Object single words. For the anterior hippocampus ROI, two participants showed similarity scores greater than 2 standard deviations away from the mean and were removed from further analysis, leaving a sample of 43 participants. For the posterior hippocampus ROI again two participants (one of whom was also excluded from the anterior hippocampus analysis) showed similarity scores greater than 2 standard deviations away from the mean and were removed from further analysis, leaving a sample of 43 participants.

For the anterior hippocampus, a repeated measures ANOVA found a significant difference between the three similarities (F_2,84_=3.40, p=0.038, η^2^=0.075). As predicted, the neural representations in the anterior hippocampus of Object word pairs were more similar to Scene single words (Figure 6A, purple bar) than to Object single words (Figure 6A, light green bar; t_42_=2.09, p=0.042, d_rm_=0.21). In fact, representations of Object word pairs were as similar to Scene single words as to themselves (Figure 6A, orange bar; t_42_=0.38, p=0.71). Object word pairs were significantly less similar to Object single words than to themselves (t_42_=2.54, p=0.015, drm=0.23). Of note, these results cannot be explained by subsequent memory performance because Scene single words and Object single words were remembered equally well (t_42_=0.68, p=0.50).

**Figure 6.**
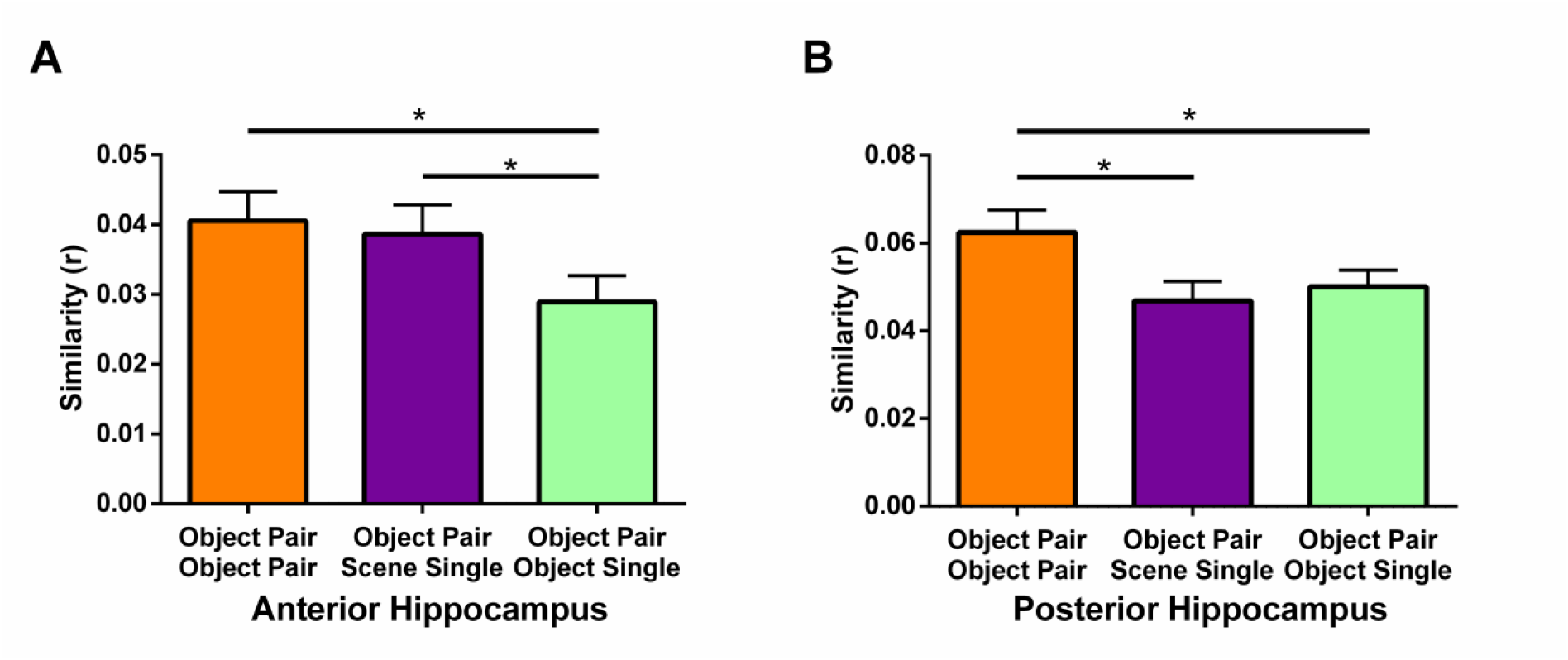
The neural similarity of Object word pairs, Scene single words and Object single words separately for the anterior and posterior hippocampus. **(A)** Anterior hippocampus. **(B)** Posterior hippocampus. Object Pair Object Pair – the similarity between Object word pairs between runs. Object Pair Scene Single – the similarity between Object word pairs and Scene single words. Object Pair Object Single – the similarity between Object word pairs and Object single words. Error bars represent 1 standard error of the mean adjusted for repeated measures (Morey, 2008). *p < 0.05.

For the posterior hippocampus, a repeated measures ANOVA also found a significant difference between the three similarities (F_2_,_84_=4.83, p=0.010, η^2^=0.10). However, in contrast to the anterior hippocampus, the neural representations in the posterior hippocampus of Object word pairs were more similar to themselves (Figure 6B, orange bar) than either Scene single words (Figure 6B, purple bar; t_42_=2.60, p=0.013, drm= 0.32) or Object single words (Figure 6B, light green bar; t_42_=2.33, p=0.025, drm=0.26). Moreover, there was no difference between the representations of Scene and Object single words (t_42_=-0.71, p=0.48). As before, these results cannot be explained by subsequent memory performance because Scene single words and Object single words were remembered equally well (t_42_=0.74, p=0.46).

Overall, these multivariate results show that within the anterior hippocampus, Object word pairs were represented in a similar manner to Scene single words, but not Object single words. On the other hand, within the posterior hippocampus, Object word pairs were only similar to themselves. This provides further support for our hypothesis that Object word pairs evoke anterior (but not posterior) hippocampal activity when scene imagery is involved.

#### VVIQ and the use of imagery

As well as examining participants in one large group, as above, we also divided them into three groups based on whether they reported high, mid or low imagery ability on the VVIQ. We found no differences in memory performance among the groups on the word pair tasks (F<0.4 for all contrasts). Similarly, fMRI univariate analyses involving the word pair conditions revealed no differences in hippocampal activity. Voxel based morphology (VBM; Ashburner, 2009; Ashburner & Friston, 2000; Mechelli, Price, Friston, & Ashburner, 2005) showed no structural differences between the groups anywhere in the brain, including in the hippocampus.

Interestingly, however, the imagery groups did differ in one specific way – their strategy for processing the Object word pairs. While strategy use was similar across the imagery groups for the other word conditions, for the Object word pairs, twice as many participants indicated using a scene imagery strategy in the high imagery group (n=12/15; 80%) than in the mid or low imagery groups (n=5/15; 33% and 6/15; 40% respectively). Comparison of scene strategy use compared to other strategy use across the imagery groups revealed this to be a significant difference (χ^2^ (2) = 7.65, p = 0.022).

Given this clear difference in scene imagery use specifically for the Object word pairs, we performed the anterior and posterior hippocampus RSA analyses again for the three imagery participant groups. We hypothesised that in the anterior hippocampus, the high imagery group would represent Object word pairs in a similar manner to Scene single words (as with our whole group analyses) whereas this would not be the case in the mid or low imagery groups. For the posterior hippocampus, on the other hand, we expected no differences between the imagery groups. Participants with similarity values greater than 2 standard deviations away from the mean were again excluded. For the anterior hippocampus ROI analyses this resulted in one participant being removed from each group. For the posterior hippocampus two participants were excluded (both different participants to those excluded from the anterior hippocampus analyses), one from the mid imagery group and one from the low imagery group. Importantly, the pattern of scene imagery strategy remained the same even after the removal of these few participants (anterior hippocampus: high imagery group, n=11/14; mid imagery group, n=5/14; low imagery group, n=5/14; χ^2^ (2) = 6.86, p = 0.032; posterior hippocampus: high imagery group, n=12/15; mid imagery group, n=4/14; low imagery group, n=5/14; χ^2^ (2) = 9.10, p = 0.011).

As predicted, in the anterior hippocampus for the high imagery group, Object word pairs were more similar to Scene single words than Object single words (Figure 7A; t_13_=4.63, p<0.001, d=0.78). This was not the case for the mid or low imagery groups (t_13_=0.472, p=0.65; t_13_=0.20, p=0.85, respectively). Of note, the interaction between the imagery groups was significant (Figure 7B; F_2,39_=3.53, p=0.039, η^2^=0.15). Independent samples t-tests showed that the difference between the similarities was greater in the high imagery group than in the mid and low imagery groups (t_26_=2.09, p=0.046, d=0.79; t_26_=2.72, p=0.011, d=1.03, respectively). As before, these differences cannot be explained by subsequent memory performance because all three groups showed no differences between the Scene single and Object single words (high imagery group: t_13_=0.35, p=0.74; mid imagery group: t_13_=0.40, p=0.69; low imagery group: t_13_=1.18, p=0.26).

**Figure 7.**
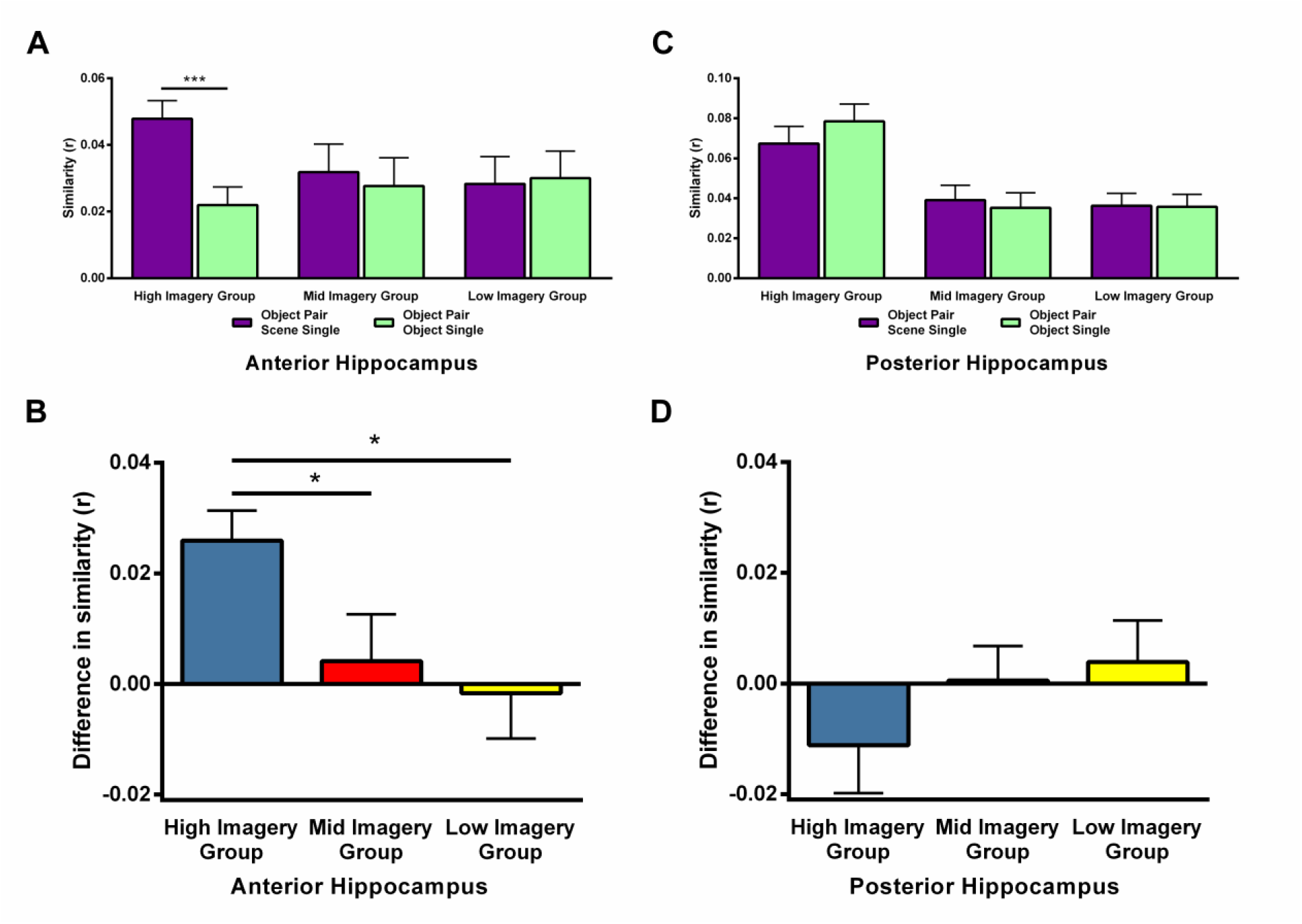
RSA comparisons of the three imagery groups separately for the anterior and posterior hippocampus. **(A)** The neural similarity of Object word pairs, Scene single words and Object single words in the anterior hippocampus when split by self-reported imagery use. Object Pair Scene Single – the similarity between Object word pairs and Scene single words. Object Pair Object Single – the similarity between Object word pairs and Object single words. **(B)** The difference in similarity between Object word pairs and Scene single words compared to Object words pairs and Object single words in the imagery groups in the anterior hippocampus. **(C)** The neural similarity of Object word pairs, Scene single words and Object single words in the posterior hippocampus when split by self-reported imagery use. Object Pair Scene Single – the similarity between Object word pairs and Scene single words. Object Pair Object Single – the similarity between Object word pairs and Object single words. **(D)** The difference in similarity between Object word pairs and Scene single words compared to Object words pairs and Object single words in the imagery groups in the posterior hippocampus. Error bars represent 1 standard error of the mean. *p < 0.05, ***p<0.001.

For the posterior hippocampus, on the other hand, there were no differences in similarities in any of the imagery groups (Figure 7C; high imagery group: t_14_=-1.29, p=0.22; mid imagery group: t_13_=0.50, p=0.63; low imagery group: t_13_=0.084, p=0.94). In line with these findings, the interaction between the imagery groups was also not significant (Figure 7D; F_2,40_=1.07, p=0.35). As before, there were no differences in subsequent memory performance between the Scene single and Object single words, suggesting this was not influencing the activity patterns (high imagery group: t_14_=0.40, p=0.69; mid imagery group: t_13_=-0.06, p=0.95; low imagery group: t_13_=1.25, p=0.24).

In summary, the neural patterns in anterior hippocampus for Object word pairs showed greater similarity with the Scene single words in the high imagery group, whereas for the mid and low imagery groups this was not the case. On the other hand, we saw no differences in any of the imagery groups in the posterior hippocampus. This provides further evidence linking the anterior hippocampus with the processing of Object word pairs through scene imagery.

## DISCUSSION

The aim of this study was to understand the role of the hippocampus in processing verbal paired associates (VPA). There were five findings. First, we observed greater anterior (but not posterior) hippocampal activity for high imagery (concrete) word pairs than very low imagery (abstract) word pairs, highlighting the influence of visual imagery. Second, very low imagery abstract word pairs compared to very low imagery abstract single words revealed no differences in hippocampal engagement, despite the former involving binding, adding further support for the significance of visual imagery. Third, increased anterior (but not posterior) hippocampal engagement was apparent for Scene word pairs more than Object word pairs, implicating specifically scene imagery. Fourth, for Object word pairs, fMRI activity patterns in the anterior (but not posterior) hippocampus were more similar to those for scene imagery than object imagery, further underlining the propensity of the anterior hippocampus to respond to scene imagery. Finally, our examination of high, mid and low imagery users found that the only difference between them was the use of scene imagery for encoding Object word pairs by high imagers, which in turn was linked to scene-related activity patterns in the anterior (but not posterior) hippocampus. Overall, our results provide evidence that anterior hippocampal engagement during VPA seems to be closely related to the use of scene imagery, even for Object word pairs.

Previous findings have hinted that visual imagery might be relevant in the hippocampal processing of verbal material such as VPA. Work in patients with right temporal lobectomies, which included removal of some hippocampal tissue, suggested that while memory for high imagery word pairs was impaired, memory for low imagery word pairs was preserved (Jones-Gotman & Milner, 1978). Furthermore, instructing these patients to use visual imagery strategies impaired both high and low imagery word pair performance (Jones-Gotman, 1979). More recently, detailed examination of the language use of patients with bilateral hippocampal damage showed that the patients used fewer high imagery words when producing verbal narratives compared to both healthy controls and patients with damage elsewhere in the brain (Hilverman et al., 2017), supporting a link between the hippocampus and word imageability. In addition, higher than expected word pair performance has been found in amnesic patients for highly semantically related word pairs in comparison to unrelated word pairs of the kind that are usually employed in VPA tasks (Shimamura & Squire, 1984; Winocur & Weiskrantz, 1976). This suggests that when alternate strategies can be used to remember word pairs (i.e. using their semantic relationship rather than constructing scene imagery) amnesic patients do not show the typical VPA impairment. We are, however, unaware of any study that has examined VPA in patients with selective bilateral hippocampal damage where high and low imagery word pairs were directly compared (Clark & Maguire, 2016).

fMRI findings also support a possible distinction in hippocampal engagement between high and low imagery word pairs. Caplan and Madan (2016) investigated the role of the hippocampus in boosting memory performance for high imagery word pairs, concluding that imageability increased hippocampal activity. However, greater hippocampal activity for high over low imagery word pairs was only observed at a lenient whole brain threshold (p<0.01 uncorrected, cluster size ≥ 5), possibly because their low imagery words (e.g., muck, fright) retained quite a degree of imageability. Furthermore, they did not examine the influence of different types of visual imagery on hippocampal engagement.

We did not find hippocampal engagement for the low imagery Abstract word pairs compared to Abstract single words, even when using ROI analyses and just the remembered stimuli. We acknowledge that null results can be difficult to interpret and that an absence of evidence is not evidence of absence. However, even our lenient uncorrected ROI analyses found no evidence of increased hippocampal activity. This is in clear contrast to the finding of increased hippocampal activity for the high imagery word pairs over the very low imagery Abstract word pairs at the whole brain level. The most parsimonious interpretation is, therefore, that Abstract word pairs may be processed differently to the high imagery word pairs, in particular in terms of hippocampal engagement.

By contrast, activity associated with the Abstract word pairs was evident outside of the hippocampus, where regions that included the left middle temporal cortex, left temporal pole and left inferior frontal gyrus were engaged. These findings are in line with other fMRI studies that examined the representations of abstract words and concepts in the human brain (Binder, Westbury, McKiernan, Possing, & Medler, 2005; Wang, Conder, Blitzer, & Shinkareva, 2010; Wang et al., 2017). Our results, therefore, align with the notion of different brain systems for processing concrete (high imagery) and abstract (low imagery) concepts and stimuli.

Our different word types were extremely well matched across a wide range of features, with the abstract words being verified as eliciting very little imagery, and the scene and object words as reliably eliciting the relevant type of imagery. Using these stimuli we showed that hippocampal involvement in VPA is not linked to visual imagery in general but seems to be specifically related to scene imagery, even when each word in a pair denoted an object. This supports a prediction made by Maguire and Mullally (2013; see also Clark & Maguire, 2016), who noted that a scene allows us to collate a lot of information in a quick, coherent and efficient manner. Consequently, they proposed that people may automatically use scene imagery during the processing of high imagery verbal material. For instance, we might visualise the scene within which a story is unfolding, or place the objects described in word pairs in a simple scene together.

If verbal tasks can provoke the use of imagery-based strategies, and if these strategies involve scenes, then patients with hippocampal amnesia would be expected to perform poorly on VPA tasks involving high imagery concrete words because they are known to have difficulty with constructing scenes in their imagination (e.g. Andelman, Hoofien, Goldberg, Aizenstein, & Neufeld, 2010; Hassabis, Kumaran, Vann, et al., 2007; Kurczek et al., 2015; Mullally, Intraub, & Maguire, 2012; Race et al., 2011). This impairment, which was not apparent for single objects, prompted the proposal of the scene construction theory which holds that scene imagery constructed by the hippocampus is a vital component of memory and other functions (Hassabis & Maguire, 2007; Maguire & Mullally, 2013). Findings over the last decade have since linked scenes to the hippocampus in relation to autobiographical memory (Hassabis, Kumaran, & Maguire, 2007; Hassabis & Maguire, 2007) but also widely across cognition, including perception (Graham et al., 2010; McCormick, Rosenthal, et al., 2017; Mullally et al., 2012), future-thinking (Hassabis, Kumaran, Vann, et al., 2007; Irish, Hodges, & Piguet, 2013; Schacter et al., 2012), spatial navigation (Clark & Maguire, 2016; Maguire, Nannery, & Spiers, 2006) and decision-making (McCormick, Rosenthal, Miller, & Maguire, 2016; Mullally & Maguire, 2014). However, as the current study was only designed to examine the role of the hippocampus in the VPA task, we do not speculate further here as to whether or not scene construction is the primary mechanism at play within the hippocampus. For more on this issue, we refer the reader to broader theoretical discussions of the scene construction theory (Clark & Maguire, 2016; Dalton & Maguire, 2017; Maguire, Intraub, & Mullally, 2016; McCormick, Ciaramelli, De Luca, & Maguire, 2018) and alternative accounts of hippocampal function (Eichenbaum & Cohen, 2014; Moscovitch et al., 2016; Schacter et al., 2012; Sheldon & Levine, 2016).

Our hippocampal findings were located in the anterior portion of the hippocampus. Anterior and posterior functional differentiation is acknowledged as a feature of the hippocampus, although the exact roles played by each portion are not widely agreed (Fanselow & Dong, 2010; Moser & Moser, 1998; Poppenk, Evensmoen, Moscovitch, & Nadel, 2013; Ritchey, Montchal, Yonelinas, & Ranganath, 2015; Strange, Witter, Lein, & Moser, 2014). Of note, the medial portion of the anterior hippocampus contains the presubiculum and parasubiculum hippocampal sub-fields. These areas have been highlighted as being consistently implicated in scene processing (reviewed in Zeidman & Maguire, 2016) and were recently proposed to be neuroanatomically determined to process scenes (Dalton & Maguire, 2017). The current results seem to accord with these findings, although higher resolution studies are required to determine the specific subfields involved.

An important point to consider is whether our results can be explained by the effectiveness of encoding, as measured in a subsequent memory test. It is certainly true that people tend to recall fewer abstract than concrete words in behavioural studies of memory (Jones, 1974; Paivio, 1969; Paivio, Walsh, & Bons, 1994). We tested memory for both single words and paired words. Memory performance for Scene, Object and Abstract words was comparable when tested singly. Memory for the word pairs was significantly lower for the low imagery Abstract word pairs compared to the Scene word pairs and Object word pairs. Nevertheless, performance for all conditions was above chance, which was impressive given the large number of stimuli to be encoded with only one exposure. Increased hippocampal activity was apparent for both Scene word pairs and Object words pairs compared to the Abstract word pairs when all stimuli or only the subsequently-remembered stimuli were analysed. Furthermore, while Object word pairs were remembered better than Scene word pairs, hippocampal activity was nevertheless greater for the Scene word pairs. This shows that our results cannot be explained by encoding success. It is also worth considering why such a gradient in memory performance was observed within the word pairs, with Object word pairs being remembered better than Scene word pairs, which were remembered better than Abstract word pairs. One possibility may be that memory performance benefitted from the extent to which the two words could be combined into some kind of relationship. This is arguably easier for two objects than two scenes, both of which are easier than for two abstract concepts.

While differences between the performance of amnesic patients and healthy participants on VPA tasks are typically observed during cued recall, in the current study we used recognition memory tests post-scanning to assess the success of encoding. This is because testing cued recall for 135 word pairs that were each seen only once is simply too difficult even for healthy participants. For example, learning just 14 (high imagery concrete) word pairs on the WMS-IV VPA task is performed over 4 learning trials. We did, however, ensure that the associative recognition memory test was challenging by constructing the lure word pairs from the single words that were presented to the participants during scanning. Thus, all words were previously seen by participants, but not all were previously seen in pairs.

Moreover, we believe that the use of a recognition memory test instead of cued recall had little impact on the patterns of brain activity we observed because brain activity was assessed during the initial presentation of the word pairs and not during memory retrieval. As participants were not told exactly how their memory would be tested after the learning phase, it might be expected that participants engaged in the most effortful encoding that they could. That the involvement of the hippocampus was identified when using all the trials in the fMRI analysis or just the subsequently remembered stimuli, also points to the use of imagery at the time of stimuli presentation as being of most relevance rather than encoding success.

There is a wealth of research linking the hippocampus with associative binding (e.g. Addis, Cheng, Roberts, & Schacter, 2011; Davachi, 2006; Eichenbaum & Cohen, 2014; Konkel & Cohen, 2009; Palombo, Hayes, Peterson, Keane, & Verfaellie, 2018; Rangel et al., 2016; Roberts et al., 2017; Schwarb et al., 2015). We do not deny this is the case, but suggest that our results provoke a reconsideration of the underlying reason for apparent associative effects. We found that the creation of associations between non-imageable Abstract word pairs did not elicit an increase in hippocampal activity compared to Abstract single words, even when only subsequently-remembered stimuli were considered. If binding per se was the reason for hippocampal involvement in our study, then this contrast should have revealed it. We suggest instead that the anterior hippocampus engages in associative binding specifically to create scene imagery, and that this relationship with scenes has been underestimated or ignored in VPA and other associative tasks despite potentially having a significant influence on hippocampal engagement.

Our participants were self-declared low, mid or high imagery users as measured by the VVIQ. They differed only in the degree of scene imagery usage, in particular during the processing of Object word pairs, with high imagers showing the greatest amount. Given that scene imagery has been implicated in functions across cognition, it might be predicted that those who are able to use scene imagery well might have more successful recall of autobiographical memories and better spatial navigation. Individual differences studies are clearly required to investigate this important issue in depth, as currently there is a dearth of such work. In the present study, increased use of scene imagery by the high imagery group did not convey a memory advantage for the Object word pairs. However, in the real world, with more complex memoranda like autobiographical memories, we predict that scene imagery would promote better memory.

In conclusion, we showed a strong link between the anterior hippocampus and processing words in a VPA task mediated through scene imagery. This offers a way to reconcile hippocampal theories that have a visuospatial bias with the processing and subsequent memory of verbal material. Moreover, we speculate that this could hint at a verbal system in humans piggy-backing on top of an evolutionarily older visual (scene) mechanism. We believe it is likely that other common verbal tests, such as story recall and list learning, which are typically highly imageable, may similarly engage scene imagery and the anterior hippocampus. Greater use of low imagery abstract verbal material would seem to be prudent in future verbal memory studies. Indeed, an obvious prediction arising from our results is that patients with selective bilateral hippocampal damage would be better at recalling abstract compared to imageable word pairs, provided care is taken to match the stimuli precisely. Our data do not speak to the issue of whether or not scene construction is the primary mechanism at play within the hippocampus, as our main interest was in examining VPA, a task closely aligned with the hippocampus. What our results show, and we believe in a compelling fashion, is that anterior hippocampal engagement during VPA seems to be best explained by the use of scene imagery.

## Acknowledgements

E.A.M. and I.A.C. are supported by a Wellcome Principal Research Fellowship to E.A.M. (101759/Z/13/Z) and the Centre by a Centre Award from Wellcome (203147/Z/16/Z). M.K. is supported by a Wellcome PhD studentship (102263/Z/13/Z) and a Samsung Scholarship. The authors report no conflicts of interest.

